# A cellular stress response induced by the CRISPR/dCas9 activation system is not heritable through cell divisions

**DOI:** 10.1101/2019.12.23.887224

**Authors:** Andrew D. Johnston, Alali Abdulrazak, Hanae Sato, Shahina B. Maqbool, Masako Suzuki, John M. Greally, Claudia A. Simões-Pires

## Abstract

The CRISPR/Cas9 system can be modified to perform ‘epigenetic editing’ by utilizing the catalytically-inactive (dead) Cas9 (dCas9) to recruit regulatory proteins to specific genomic locations. In prior studies, epigenetic editing with multimers of the transactivator VP16 and guide RNAs (gRNAs) was found to cause adverse cellular responses. These side effects may confound studies inducing new cellular properties, especially if the cellular responses are maintained through cell divisions - an epigenetic regulatory property. Here we show how distinct components of this CRISPR/dCas9 activation system, particularly untargeted gRNAs, upregulate genes associated with transcriptional stress, defense response, and regulation of cell death. Our results highlight a previously undetected acute stress response to CRISPR/dCas9 components in human cells, which is transient and not maintained through cell divisions.

## INTRODUCTION

The prokaryotic clustered regularly interspaced short palindromic repeats (CRISPR) system has been extensively used for eukaryotic genome editing, allowing precise point mutations, insertions and deletions, as well as epigenetic editing.^1–3^ Tempering the the promise of CRISPR/Cas9 systems is the concern of off-target effects. The Cas9 nuclease protein has been shown to bind promiscuously across the genome,^4^ resulting in undesirable insertion-deletion events as a consequence of this off-target cleavage.^5^

Epigenetic editing uses dCas9 (dead Cas9), a mutated Cas9 devoid of endonuclease activity, allowing the recruitment of effector proteins to specific loci without causing mutations at those sites. Over time, different CRISPR activation (epigenetic editing) systems have been proposed and compared in regards to their efficacy and off-target effects.^6^ The first constructs consisted of the standard activator VP64 (four copies of VP16) linked to the C-terminus of dCas9.^7, 8^ VP16 is a viral protein that forms a transcriptional regulatory complex in host cells to induce early gene transcription upon herpes simplex infection.^9^ Subsequent CRISPR activation systems have been developed, many of them expressing VP16 repeats (VP64 or VP160), either fused to dCas9^7, 10–13^ or recruited by protein tagging and programmable RNA scaffolds.^14, 15^ Off-target activation has not been detected using CRISPR activation, suggesting that guide RNA (gRNA) sequences are not inducing off-target recruitment of dCas9 leading to gene activation. However, a prior study points to a possible side effect of epigenetic editing using VP64 that involves the downregulation of the Interleukin 32 gene (*IL32*).^7^ Moreover, when produced via *in vitro* transcription, CRISPR gRNAs triggered side effects related to the innate immune response in human cells, with the upregulation of genes involved in the type I interferon response.^16, 17^

Given these potential side effects of epigenetic editing, we aimed to investigate the genome-wide, off-target effects of the CRISPR components on human transcriptional regulation. Here we examined the gene expression effects of distinct components of a VP16-based CRISPR/dCas9 activation system, by analyzing cells transiently transfected with different combinations of dCas9, gRNAs and VP16 repeats, applying normalized transfected DNA amounts, and selection of positively transfected cells. This strategy allowed us to characterize a previously undetected acute stress response to the CRISPR/dCas9 components in human cells.

## MATERIAL AND METHODS

### Plasmid construction

To generate the dCas9 vectors, plasmid pAC154-dual-dCas9VP160-sgExpression (Addgene plasmid # 48240)^18^ was linearized to introduce the 2A-GFP sequence downstream to the dCas9-VP160 fusion. Reverse complement oligonucleotides were annealed and amplified to generate the 2A sequence. The GFP sequence was amplified by PCR from plasmid pBI-MCS-EGFP (Addgene plasmid #16542)^19^ and all fragments were Gibson assembled to provide the sgRNA-dCas9-VP160-2A-GFP vector. Additional steps of plasmid digestion, gel purification, and Gibson assembly were then applied to the resulting vector. In this way, distinct CRISPR components were sequentially removed to generate the vectors sgRNA-dCas9-2A-GFP and sgRNA-2A-GFP.

A gRNA cloning vector (Addgene plasmid #41824) was used as the gRNA empty backbone and for cloning the gRNA sequences as previously described.^20^ The vector was linearized, then reverse complement oligonucleotides containing the 19-nucleotide gRNA target sequence and the gRNA scaffold were annealed and Gibson assembled into the vector to generate individual gRNAs1-6. The gRNA sequences (**Supplementary Table 1**) were selected as those with the highest scores and shortest distance to the TSS using the CRISPR design tool crispr.mit.edu. Plasmid sequences are provided in **Supplementary File 1**.

### Cell culture, transfection, and sorting

HEK 293T cells were cultured in DMEM medium, supplemented with 10% fetal bovine serum (FBS, Benchmarck), 100 units/mL penicillin, and 100 μg/mL streptomycin (Life Technologies). Cells were cultured in 75 cm^2^ tissue culture flasks (NUNC, Thermo Scientific) at 37°C in a 5% CO_2_ incubator. For each condition, a total of 10^6^ cells/100 mm dish was cultured in triplicate overnight, then transfected with 1.93 pmol of GFP-expressing vectors and 3.47 pmol of gRNA vectors (**Supplementary Table 2**). Control cells received transfection reagents only. Transfections were conducted with Lipofectamine 2000 (Invitrogen) according to the manufacturer’s instructions. After 24 h following transfection, the medium was replaced and cells were kept under culture for a total time of 48 h after transfection. Subsequently, cells were detached with EDTA, pelleted, washed twice, and resuspended in FACS buffer (Hank’s balanced salt solution buffer supplemented with 1% BSA and 0.5 mM EDTA). Cell suspensions were then submitted to cell analysis and sorting in a FACSAria II cytometer (BD Biosciences). FACS data were analyzed using FACSDiva software (Becton Dickinson) with gating of single cells using FSC/W and SSC/W, and gating of GFP+ cells. When subsequent analyses were to be performed, cells were sorted into culture medium, washed twice with PBS, and pelleted.

### CD34 FACS analysis

Cells were detached with EDTA, washed twice, and suspended in FACS buffer at 5 x 10^5^ cells/mL. For each sample, three aliquots of 100 μL were prepared to be treated with CD34 PE monoclonal antibody (clone 4H11, eBioscience), isotype control PE Mouse IgG1 kappa (clone P3.6.2.8.1), and FACS buffer, respectively. Each aliquot was first treated with 20 μL of Fc receptor binding for 10 min on ice, then with 5 μL of PE antibody or buffer for 20 min on ice. After incubation, cells were washed (2 x 1 mL) and suspended in 500 μL of FACS buffer. FACS data were analyzed using FACSDiva (Becton Dickinson) or FloJow v10.5 (FlowJo LLC) software, with gating of single cells using FSC/W and SSC/W, and gating of GFP+ and CD34 PE+ cells.

### Total RNA extraction and quantitative reverse-transcription polymerase chain reaction (qRT-PCR)

Cell pellets were treated with QIAzol lysis reagent (Qiagen) and total RNA was isolated using the miRNAeasy kit (Quiagen) combined with DNAse (Qiagen) treatment according to manufacturer’s instructions. Synthesis of cDNA was performed with SuperScript III First-Strand Synthesis System for RT-PCR (Life technologies) using random hexamers as primers. *CD34*, *DDIT3*, *RELB*, and *JUNB* levels were measured with specific forward and reverse primers (**Supplementary Table 3**) with Light Cycler 480 Syber Green Master mix, according to the manufacturer’s instructions.

### RNA-seq library preparation and analysis

RNA-seq libraries were prepared from 1 ng of total RNA using the SMART-Seq HT Kit (Takara) combined with Nextera XT kit (Illumina), according to manufacturers’ instructions. One-step cDNA synthesis and double-stranded cDNA amplification was conducted with 3’ SMART-Seq CDS Primer II A for priming, and SMART-Seq HT oligonucleotide for template switching at the 5’ end of the transcript. The cDNA was then purified with the Agencourt AMPure XP kit, tagmented, and PCR amplified with appropriate index primers. Directional RNA-seq libraries were then sequenced 100 bp single-end on the Illumina HiSeq 2500. Reads were trimmed by Trim Galore (http://www.bioinformatics.babraham.ac.uk/projects/trim_galore/; v0.3.7) and then aligned to the hg38 reference genome using STAR v2.6.0c.^21^ Differentially expressed protein-coding genes were determined by applying a threshold of log_2_-fold change > 1, and FDR < 0.05, using DESeq2 v1.16.1^22^ on protein-coding gene counts normalized by housekeeping genes^23^ as input to the RUVg command within RUVseq v1.10.0.^24^ A full description of the analysis can be found on our GitHub server: https://github.com/GreallyLab/Johnston_Simoes-Pires_et_al_2019.

### Analysis of gene ontology enrichment and protein-protein associations

The list of 97 overlapping dysregulated genes was evaluated through functional enrichment analysis with DAVID (**Supplementary File 2**).^25^ A total of 30 genes from enriched pathways showing a p-value < 0.005 were further analyzed for their predicted protein associations in the STRING database.^26^

### Analysis of off-target effects

Predicted gRNA off-target sites were obtained from the CRISPOR website (http://crispor.tefor.net/crispor.py?batchId=0xd7m55fmDlcoF8EzTa9#s343+).^27^ These regions were then intersected by +/− 1 kb from TSSs of the 97 overlapping dysregulated genes using *bedtools2* v2.26.0.

### Determination of number of cell divisions

A total of 5 x 10^4^ cells, either GFP+ (CRISPR CD34) or GFP− (control) were directly sorted into wells of a 24-well plate in culture medium. Cells were cultured and passaged every 48 h until GFP+ cells turned negative under the microscope. The total number of cells were counted at every passage, and the number of cell divisions was calculated as the population doubling level (PDL) with the formula n = 3.32 (log N_24h_ – log N_0_), where N_24h_ is the total number of cells after 24 h in culture, and N_0_ is the number of cells seeded in the previous passage.

## RESULTS

To investigate the VP16-based CRISPR activation system, we first designed a vector for the human expression of both a scrambled gRNA and the dCas9 fused to ten repeats of VP16 (VP160). In order to discriminate between transfected and non-transfected cells, green fluorescent protein (GFP) was fused to the VP160 open reading frame using a linker encoding the cleavable peptide 2A.^28^ We used the system to target the endogenous activation of *CD34*, a gene which is not expressed in HEK 293 cells (https://www.proteinatlas.org).^29^ *CD34* encodes a transmembrane protein, allowing us to discriminate easily by antibody recognition the cells expressing the protein in living cells. In a prior study, VP16 repeats directly fused to dCas9 required a pool of gRNAs for robust activation,^8^ increasing the number of possible mismatches that could lead to off-target activation genome-wide. To test the off-target effects from multiple gRNA sequences, dCas9-VP160-2A-GFP was transfected in combination with six pooled gRNAs targeting the *CD34* promoter (**Fig. 1A**). Performing fluorescence-activated cell sorting (FACS), we demonstrated successfully induced endogenous expression of *CD34* in HEK 293T cells (**Fig. 1B**), with GFP+/CD34+ cells, showing an 80-fold increase in *CD34* mRNA levels (**Fig. 1C**). Interestingly, successfully transfected cells not expressing CD34 on the cell surface (GFP+/CD34-) also had an increase in *CD34* mRNA levels (**Fig. 1C**), suggesting a cell subpopulation with either delayed protein translation or a lack of membrane translocation. While the pooled gRNAs were indeed more effective in inducing CD34 trans-membrane expression compared to individual gRNAs, individual gRNA sequences seeding within a short distance (up to 100 nucleotides) from the transcriptional start site (TSS) were also successful, with expression levels increasing with decreasing distances from the TSS (**Supplementary Fig. 1**).

**Fig. 1.**
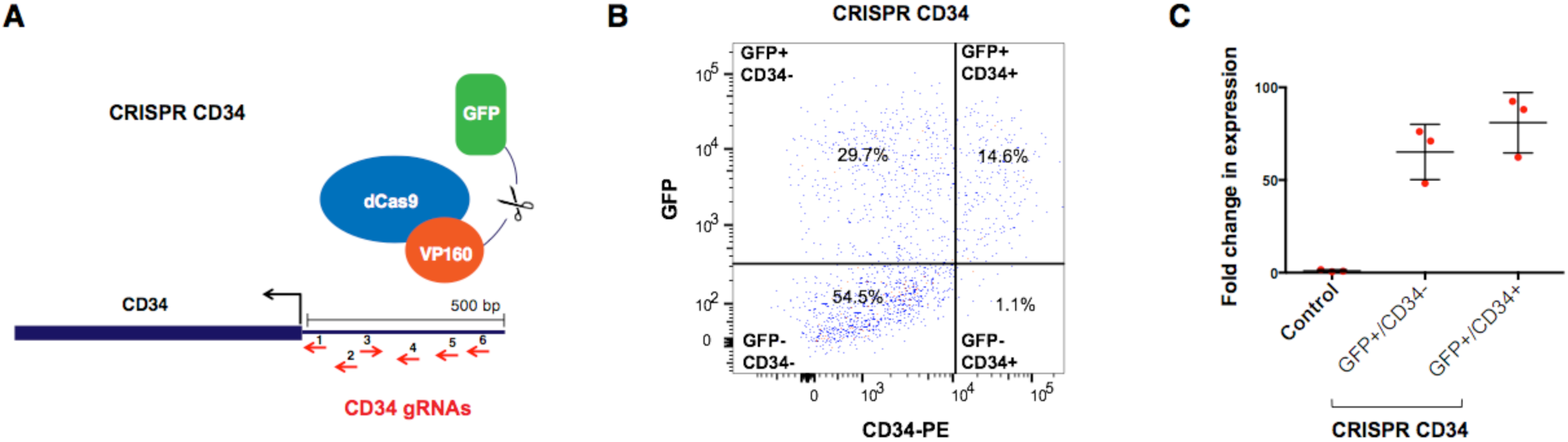
Ectopic gene activation of *CD34* with the CRISPR activation system using the sgRNA-dCas9-VP160-2A-GFP vector combined with 6 gRNAs. **A)** Overview of the CRISPR CD34 activation system: 6 gRNAs targeting the promoter of *CD34* within 500 bp from the TSS were co-transfected with a vector expressing dCas9 fused to 10 repeats of the transactivation peptide VP16, released from GFP by a cleavable peptide. **B)** FACS analysis of HEK 293 cells transfected with the CRISPR CD34 activation system. GFP+/CD34− and GFP+/CD34+ cells were sorted for subsequent analysis. **C)** The fold changes in expression in sorted cells relative to control measured by qRT-PCR.

To evaluate whether the system induced undesirable effects genome-wide, we conducted RNA-seq analyses on the GFP+ cells transfected with the full activation system including the six gRNAs (CRISPR CD34), in comparison to non-transfected cells (Control). In addition to the strong upregulation of *CD34*, a total of 161 differentially expressed genes were identified (**Fig. 2A**). We then generated a CRISPR control by sorting GFP+ cells expressing dCas9-VP160 and a scrambled gRNA (CRISPR). In this control, we detected 125 differentially expressed genes (**Fig. 2B**), with 97 of them overlapping the genes identified in the CRISPR CD34 sample (**Fig. 2C, Supplementary file 2**). Predicted gRNA off-target loci were not within 1kb of the dysregulated genes’ TSSs, suggesting that their differential expression was not a result of targeted dCas9-VP160 activation.

**Fig. 2.**
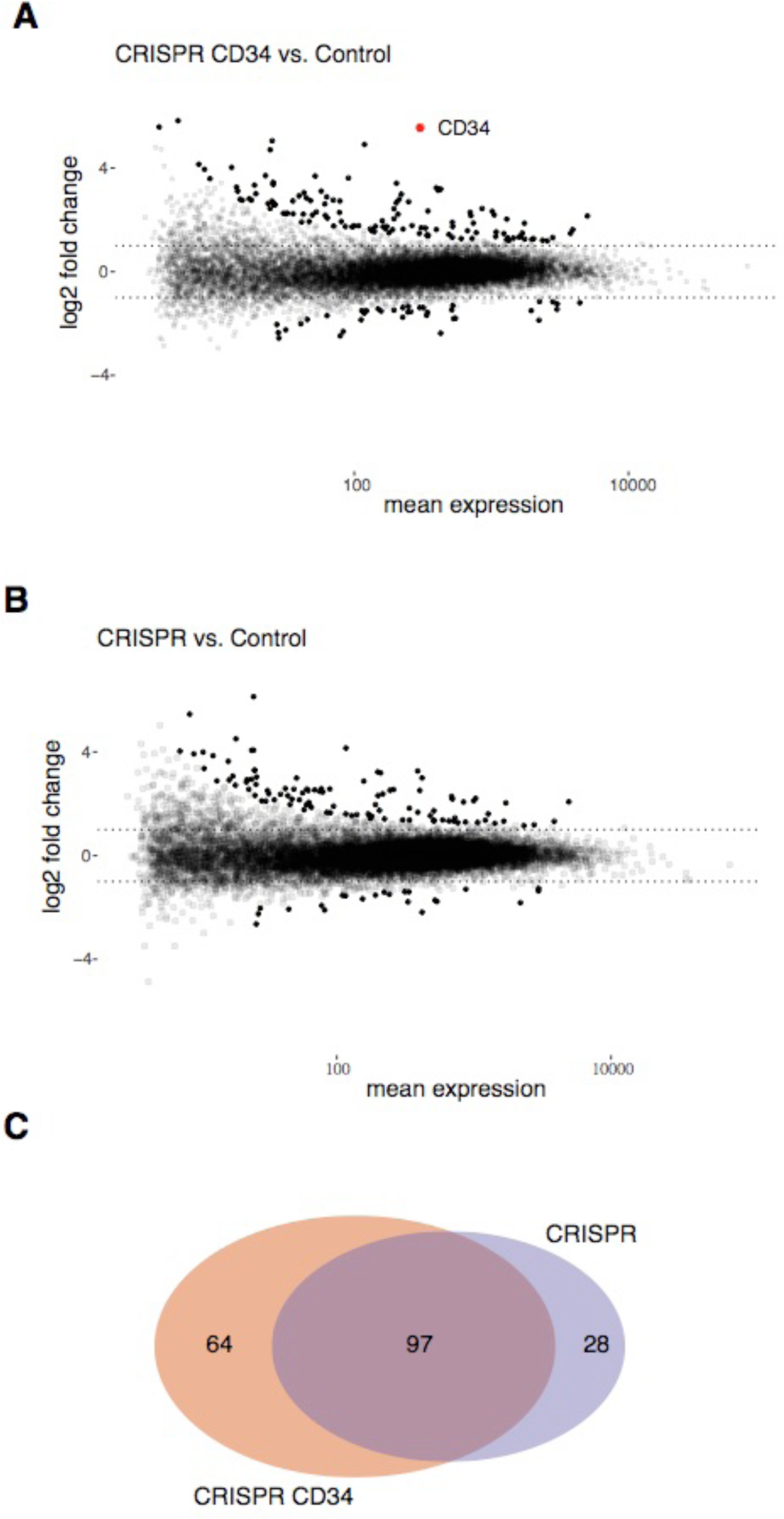
Differentially expressed genes with the CRISPR CD34 activation system and the CRISPR system in the absence of targeted gRNAs. **A)** RNA-seq MA plot of CRISPR CD34 compared with control. Black solid dots are the differentially expressed genes (log_2_ fold change > 1, FDR< 0.5). Differentially expressed *CD34* is represented by a solid red dot. **B)** RNA-seq MA plot of CRISPR control compared with control. Black solid dots are the differentially expressed genes (log_2_ fold change > 2, FDR< 0.5A). *CD34* is not differentially expressed. **C)** Venn diagram representing the 97 differentially expressed genes in common between CRISPR CD34 and CRISPR control.

Nevertheless, the consistently dysregulated genes observed in the CRISPR control cells raised the question of whether side effects may occur due to the expression of dCas9, VP16 repeats, or gRNAs. We evaluated these 97 genes through functional enrichment analysis and protein associations. The gene ontology analysis was significantly enriched for biological pathways related to apoptosis, response to cytokines, mechanical stimulus, inflammation, and response to endoplasmic reticulum stress and unfolded proteins, represented by a total of 30 genes. Further analysis of protein-protein associations related to those genes featured the pathways of cell defense and regulation of cell death (**Fig. 3**), from which we selected three node genes (*DDIT3*, *RELB*, and *JUNB*) for further investigation.

**Fig. 3.**
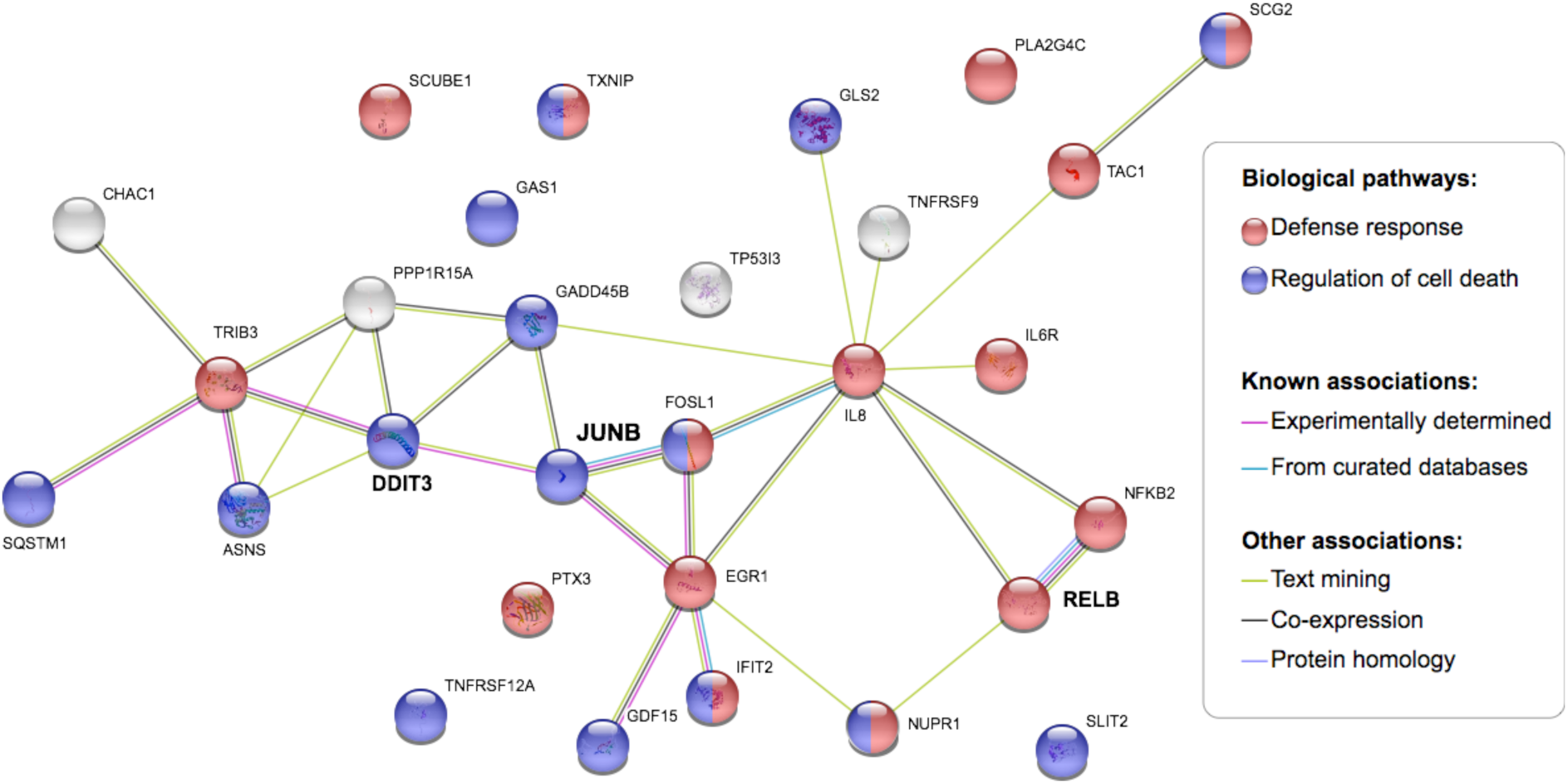
Protein-protein associations among genes selected from gene ontology analysis. Analysis from STRING database (https://string-db.org/). The genes *DDIT3*, *JUNB* and *RELB* were selected for further studies as central to the regulation of these defense response and cell death regulatory pathways.

*DDIT3* encodes the DNA Damage Inducible Transcript 3 transcription factor activated during endoplasmic reticulum stress.^30^ *RELB* is a subunit of the pleiotropic transcription factor NFκB that has a central role in cell differentiation, growth, apoptosis, inflammation, and immunity.^31–33^ *JUNB*, a component of the AP1 transcription factor, has a role in stress response and is associated with the NFκB pathway.^34–37^

Assessing the impact of the various CRISPR activation system components, we quantified the changes in expression of the selected genes in GFP+ cells transfected with distinct CRISPR components (**Fig. 4A**). Considering that the absolute amounts of foreign DNA introduced into cells may contribute to the degree of the observed stress response, we used equimolar plasmid concentrations across test conditions. First, we confirmed the activation of *CD34* in the CRISPR CD34 cells only in the presence of the targeted gRNAs; it was not induced by the expression of gRNAs alone (gRNA control) nor any other isolated component of the system (**Fig. 4B**). The stress-related genes *DDIT3*, *RELB* and *JUNB* were induced across all samples containing the CRISPR components. Expression of gRNAs in their untargeted form, either in the absence of dCas9 (gRNA control) or with a scrambled sequence in the presence of dCas9 (CRISPR and dCas9 controls), demonstrated a robust elevation of the stress-related genes’ expression (**Fig. 4B**).

**Fig 4.**
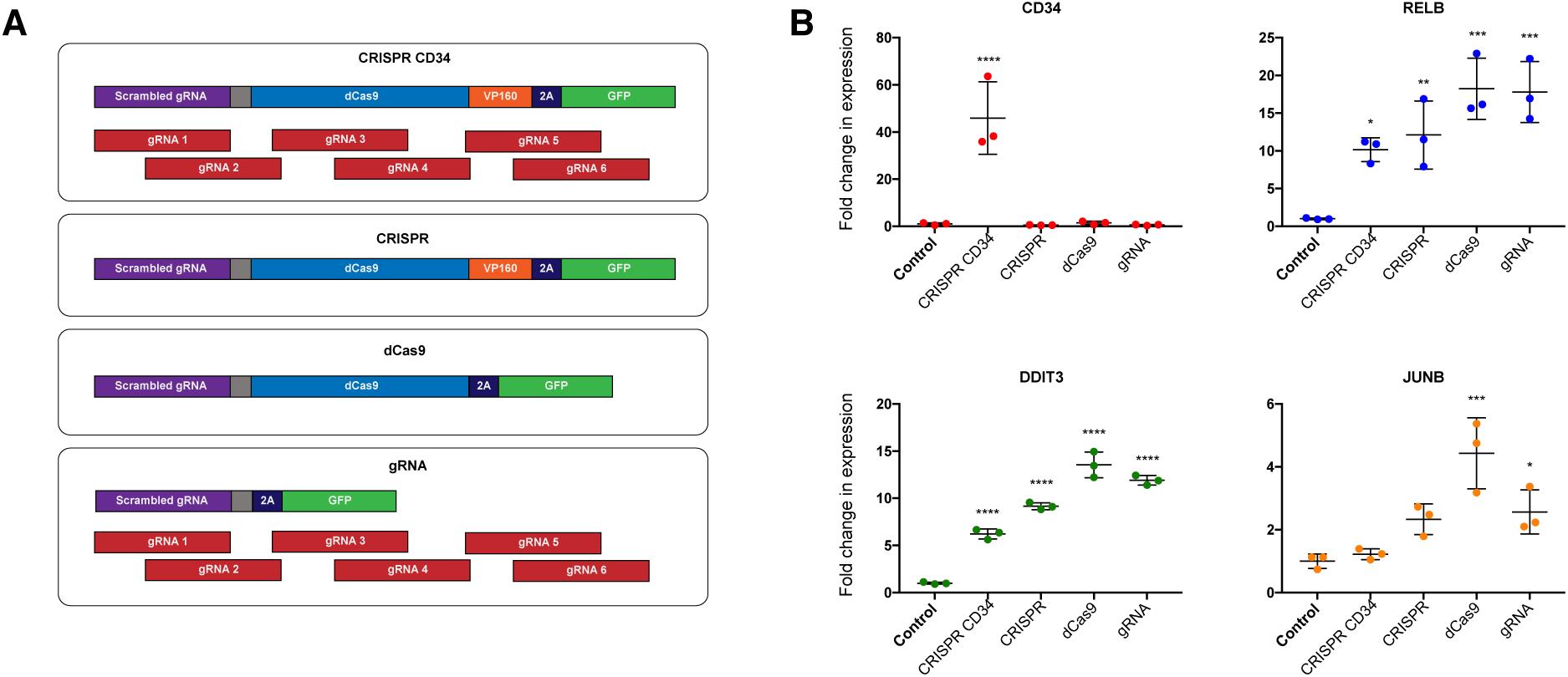
**Relative RNA expression of *CD34* and stress-related genes across CRISPR conditions**. **A)** s of the expression vectors transfected in each condition. **B)** Acute fold change in gene expression relative to control at 48 hours after transfection. P-values: *≤ 0.05; ***≤ 0.001; ****≤ 0.0001.

We then investigated whether cells transfected with the CRISPR activation system were able to return to their basal expression levels over multiple cell divisions. To do this, we kept the activated GFP+ cells in culture until cells were negative for GFP fluorescence under the microscope (after 10 cell divisions). At this point, the cells were analyzed by FACS and sorted for GFP-populations to ensure that the CRISPR components had been eliminated from the cells. We demonstrated that the upregulated stress-response genes returned to their basal levels (**Fig. 5**), indicating the absence of a memory effect for both *CD34* and the cellular stress response genes.

**Fig. 5.**
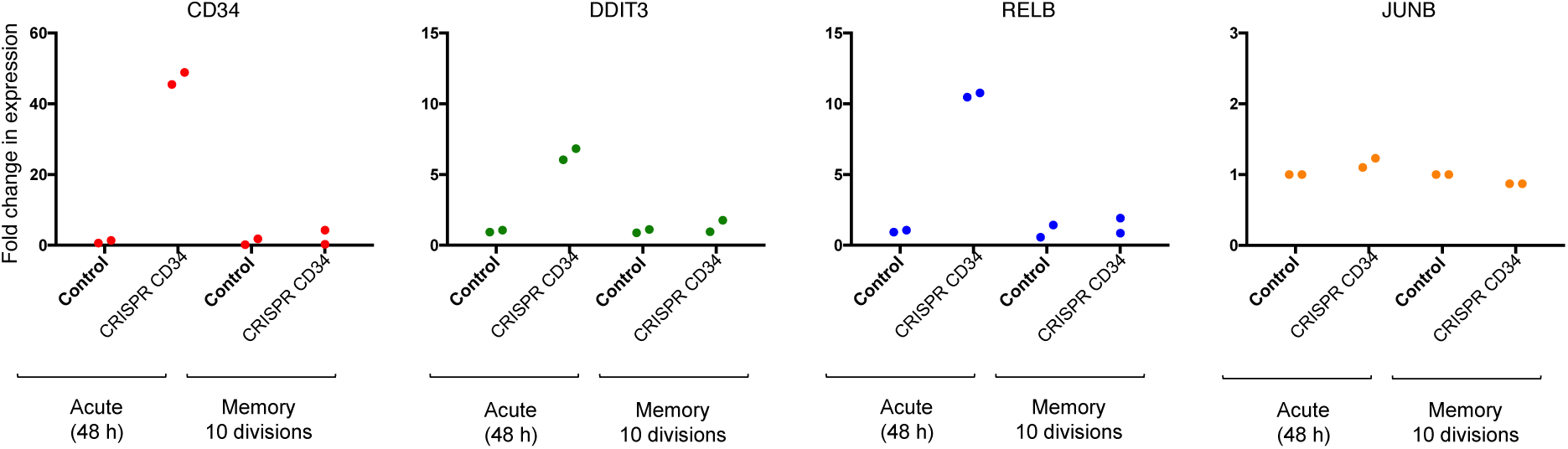
Change in gene expression relative to control in transfected cells after 10 cell divisions in comparison to the acute response. The expected induction of gene expression is seen acutely at 48 h, with complete resolution when 10 cell divisions have occurred in these GFP-cells.

## DISCUSSION

Taken together, our results point to the activation of stress genes as a side effect upon the expression of CRISPR components, especially untargeted gRNAs, not necessarily related to the presence of VP16 or to gRNA off-target sequences. Indeed, previous findings have shown that dCas9 has a higher residence time at a targeted genomic locus than at off-target loci,^38^ potentially contributing to the high specificity of gRNAs in the dCas9-VP16-based epigenetic activation systems.

The outcome of undesirable transcriptional regulation is of concern when using dCas9 fused to effectors for epigenetic editing. The changes in cellular properties resulting from epigenetic editing might be expected to be heritable, as this is one definition of cellular epigenetic properties.^39^ If side effects affecting gene expression are maintained through cell division, they will be difficult to uncouple from the desired effect of the epigenetic editing. Moreover, heritable side effects may constitute a pitfall in developing CRISPR technologies for the development of therapeutic applications.

Our findings reveal an acute cellular response to the components of the CRISPR activation system, which dissipates over the course of multiple cell divisions. While this is reassuring for the use of CRISPR-mediated epigenetic editing, we note that the effects observed involve the transient activation of transcription factors. Transient upregulation of transcription factors may induce downstream pathways, which in turn can be irreversible. One example is the role of pioneer transcription factors in somatic cell reprogramming.^40^ Accordingly, the transcription factor DDIT3, predominantly related to the stress response, has been identified as a regulatory node in erythroid lineage cell programming.^41^ Furthermore, we only examined the genome-wide expression consequences of a transient CRISPR transfection in one cell line; the potential long-term transcriptional effects of stably transfected CRISPR machinery or differing cellular response by other cell types warrant further investigation.

## CONCLUSION

An acute stress response occurs in cells when CRISPR components are used for gene activation. Although transient, the response was mediated through the upregulation of transcription factors that may, in certain cell systems, independently lead to reprogramming effects. Therefore, the impact of CRISPR components on transcription factors should be carefully taken into consideration when designing CRISPR genetic and epigenetic editing tools.

## AKNOWLEDGEMENTS

The current project has received funding from the European Union’s Horizon 2020 research and innovation programme under the Marie Sklodowska-Curie grant agreement No 750190.

## DATA AVAILABILITY

All genome sequencing data are available from the NCBI Gene Expression Omnibus database under accession number GSE11827 (https://www.ncbi.nlm.nih.gov/geo/query/acc.cgi?acc=GSE118277; reviewer token: ujijuqqqzlkjbyt).

## CODE AVAILABILITY

The code files for the all analyses are available at https://github.com/GreallyLab/Johnston_Simoes-Pires_et_al_2019.

---

**Supplementary Fig. 1.**
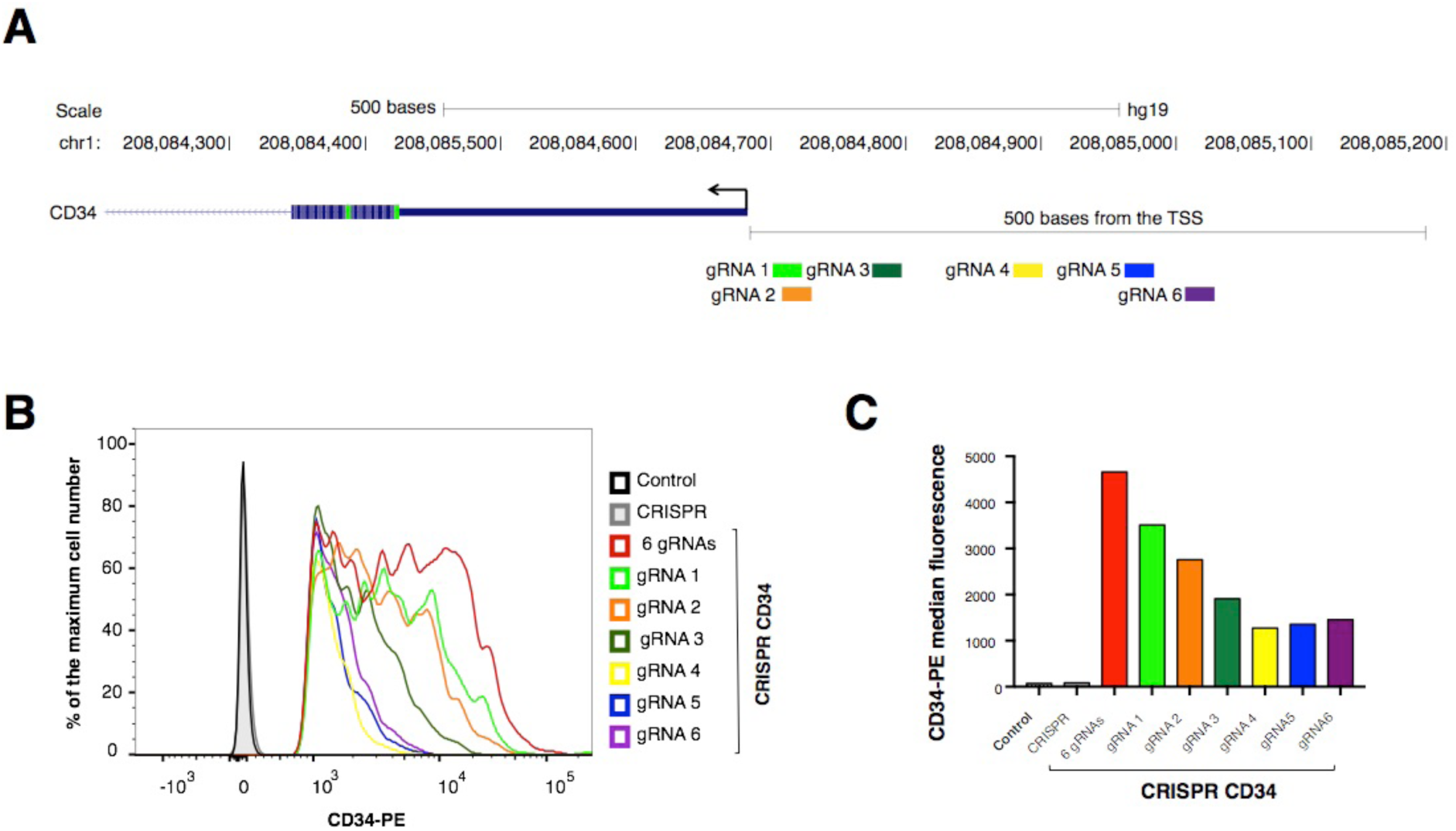
The efficiency of gRNA sequences in the activation of *CD34* using the VP16-based CRISPR activation system. **A)** The position of gRNAs 1-6 relative to the *CD34* transcriptional start site (TSS). **B)** FACS histograms depicting CD34 expression in cells transfected with the CRISPR activation system, followed by **C)** a bar graph depicting their median CD34-PE fluorescence.

**Supplementary Table 1.**
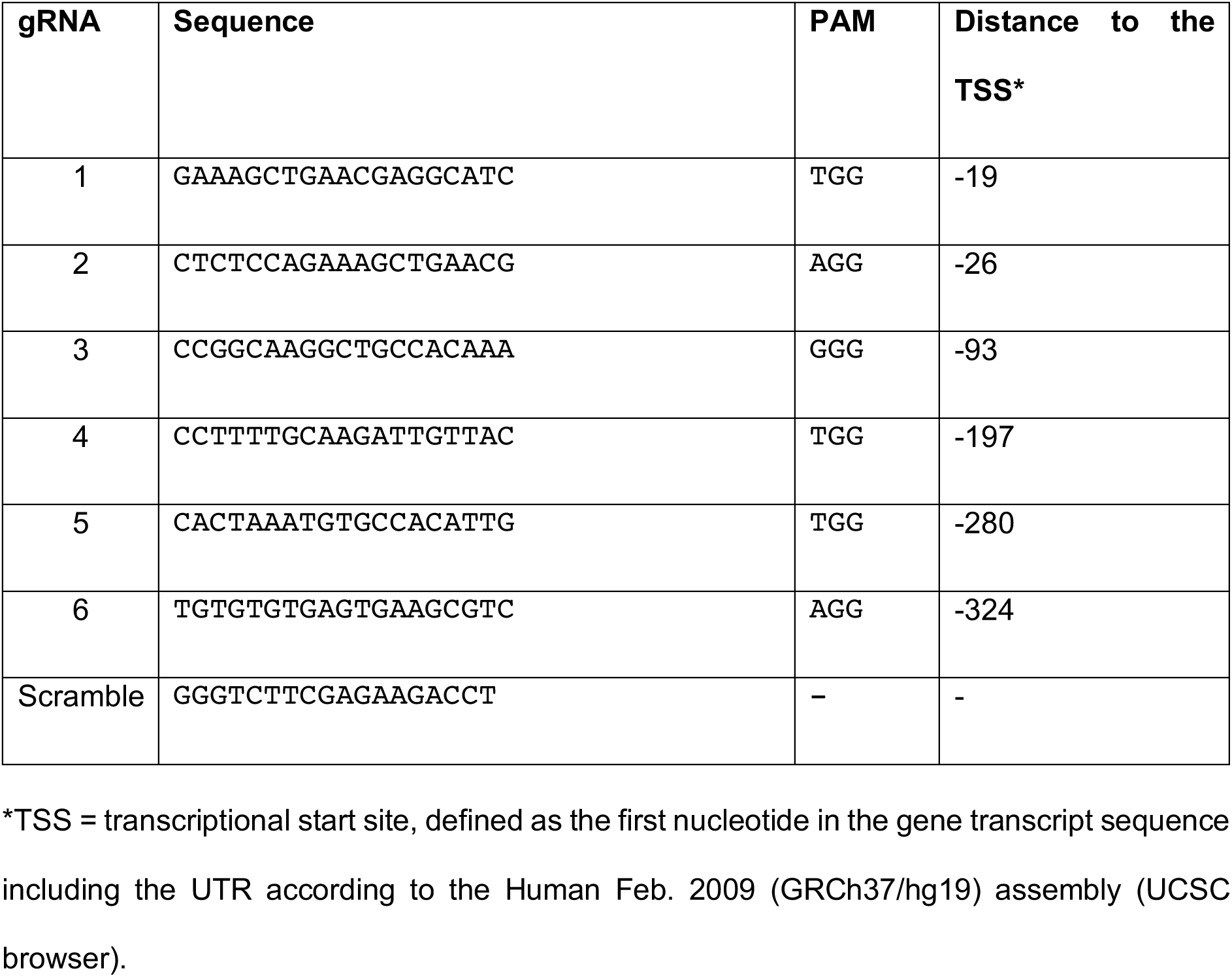
gRNA sequences.

**Supplementary Table 2.** Amount of transfected vectors per 100 mm dishes across CRISPR conditions.

| Condition | Vector (pmol) |  |  |  |  |
| --- | --- | --- | --- | --- | --- |
|  | sgRNA-dCas9-VP160-2A-GFP | sgRNA-dCas9-2A-GFP | sgRNA-2A-GFP | 6 gRNAs | gRNA cloning vector<br>(empty backbone) |
| CRISPR CD34 | 1.93 | - | - | 3.47* | - |
| CRISPR | 1.93 | - | - | - | 3.47 |
| dCas9 | - | 1.93 | - | - | 3.47 |
| gRNA | - | - | 1.93 | 3.47* | - |

\* Divided into equal amounts of each vector.

**Supplementary Table 3.**
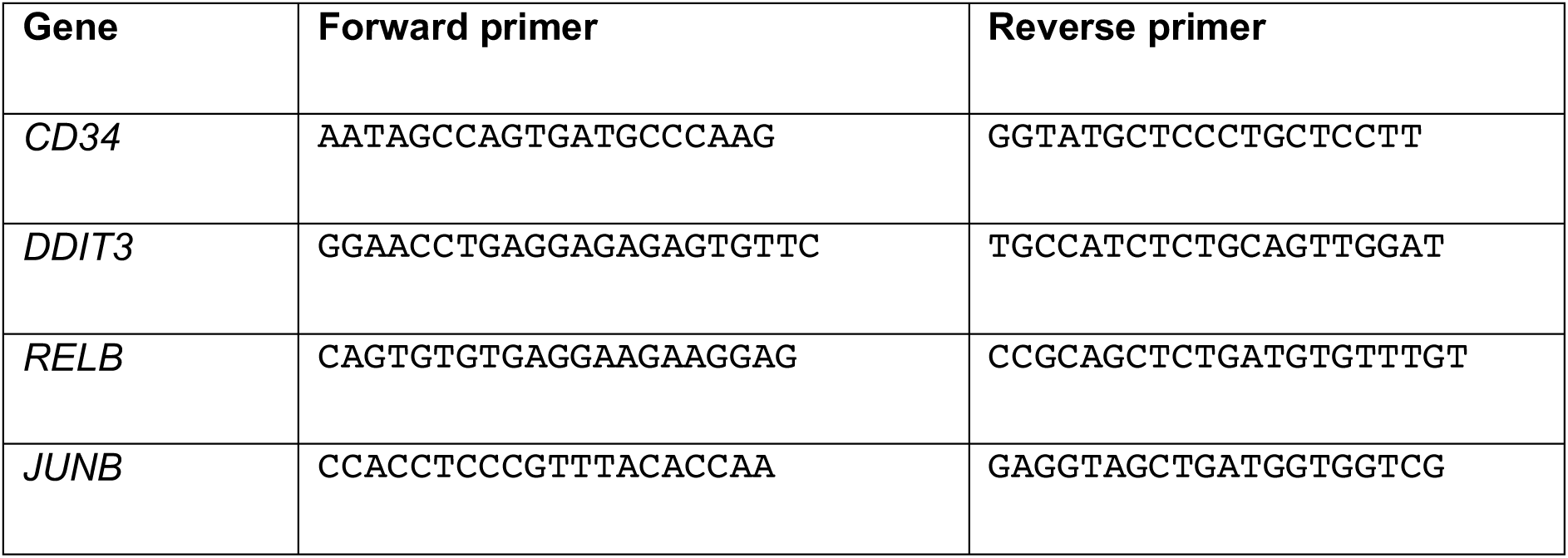
qRT PCR primers.

## Supplementary file 1

### Sequence of plasmid sgRNA-dCas9-VP160-2A-GFP

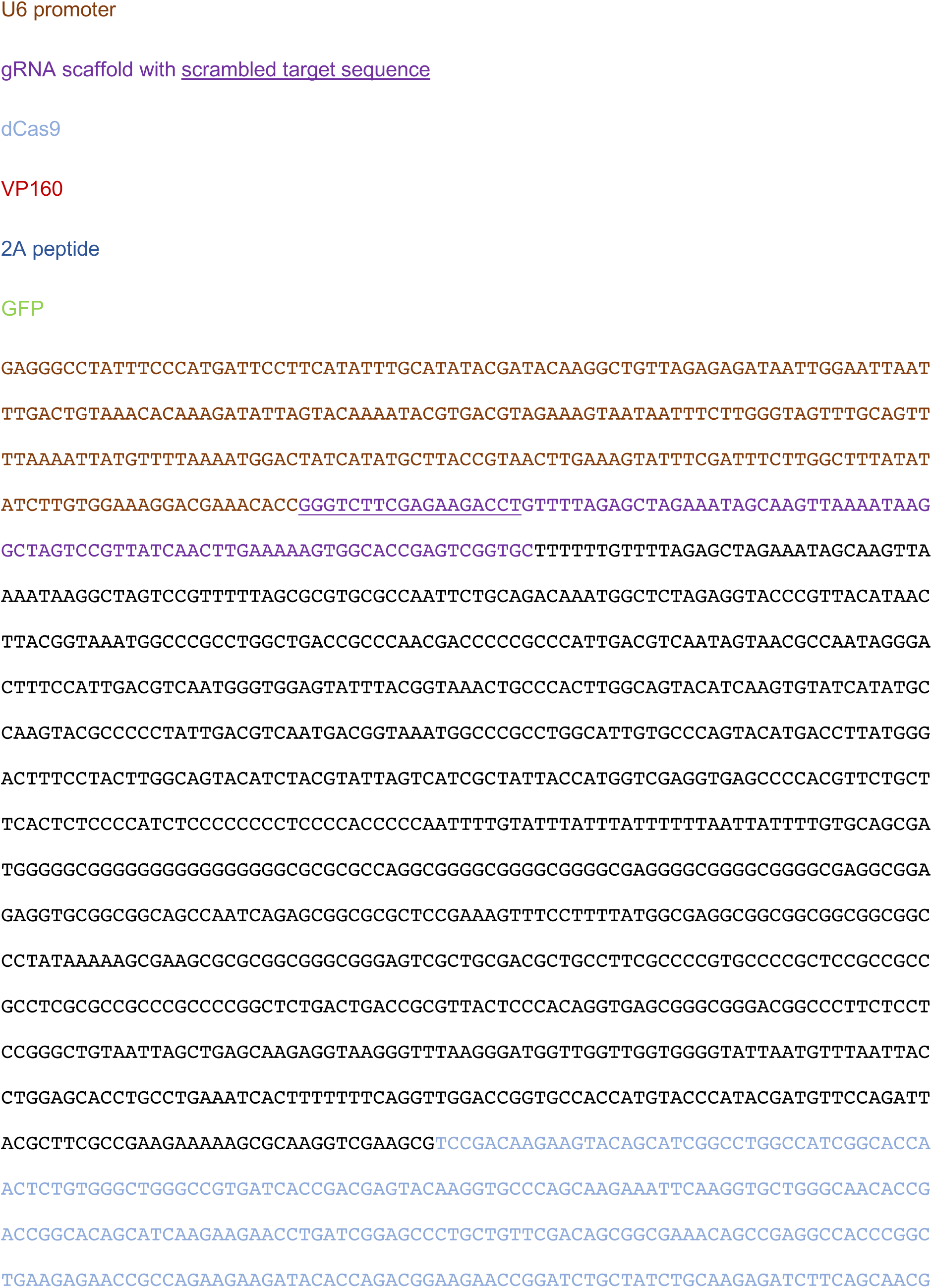
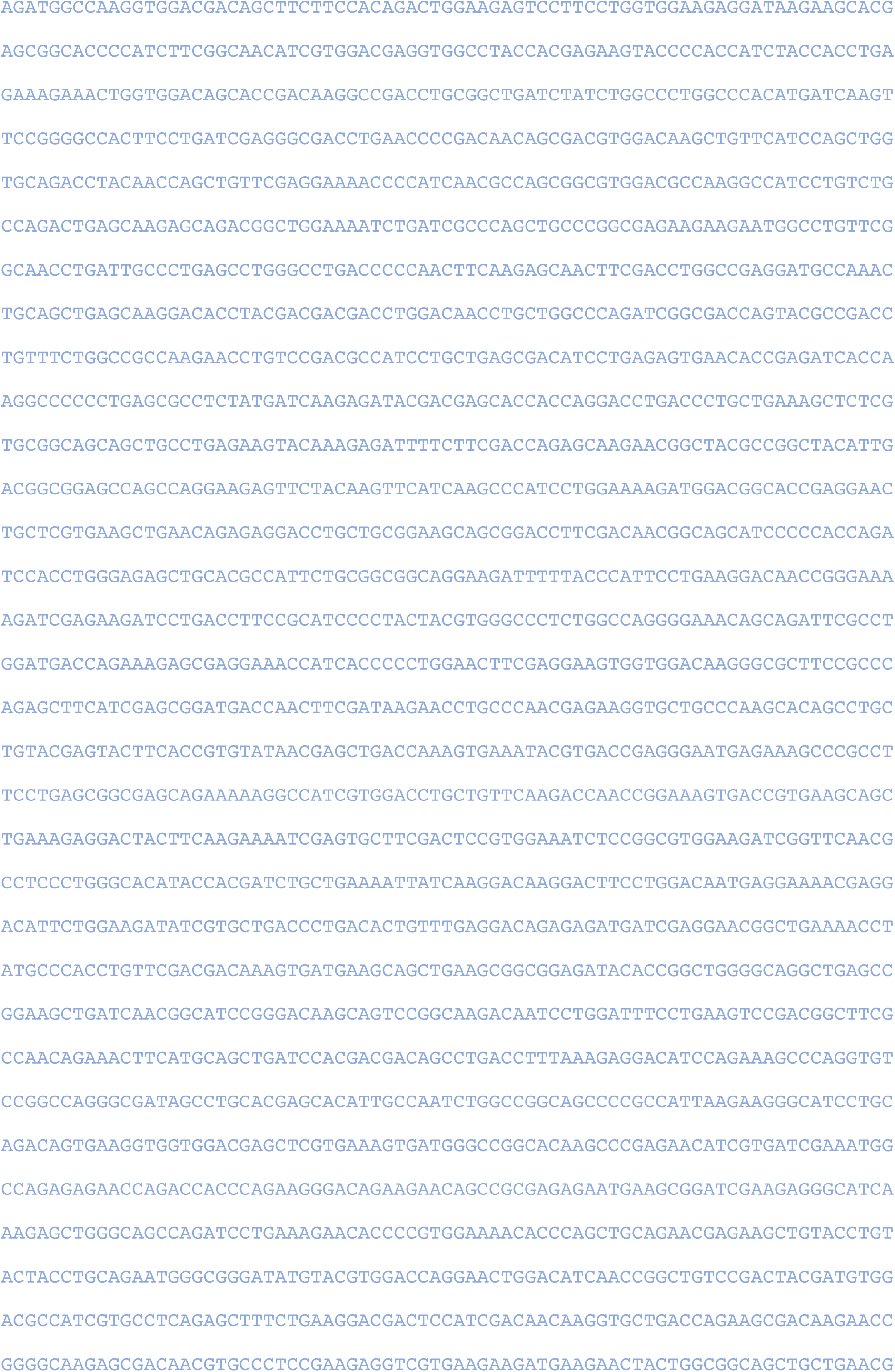
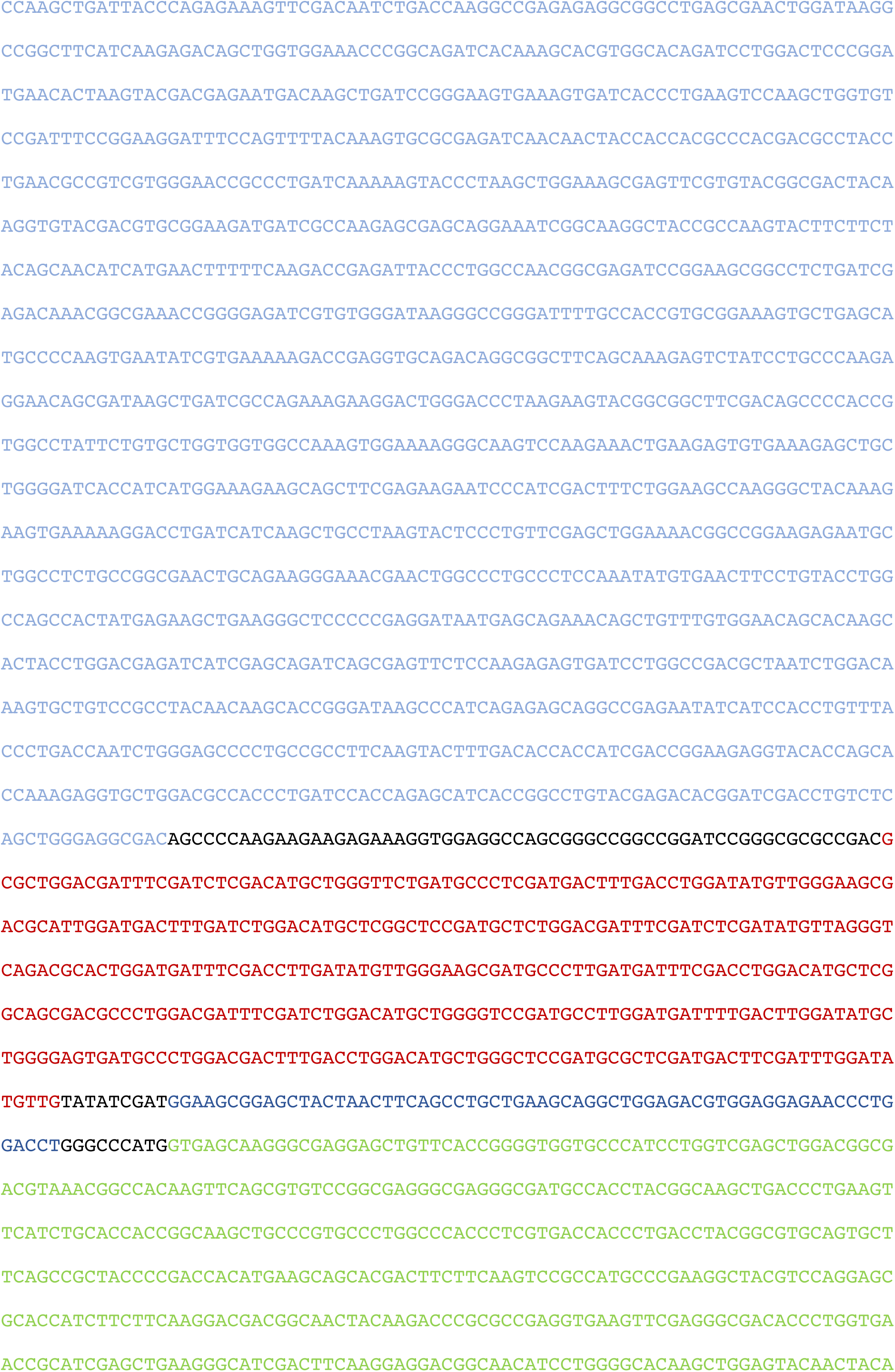
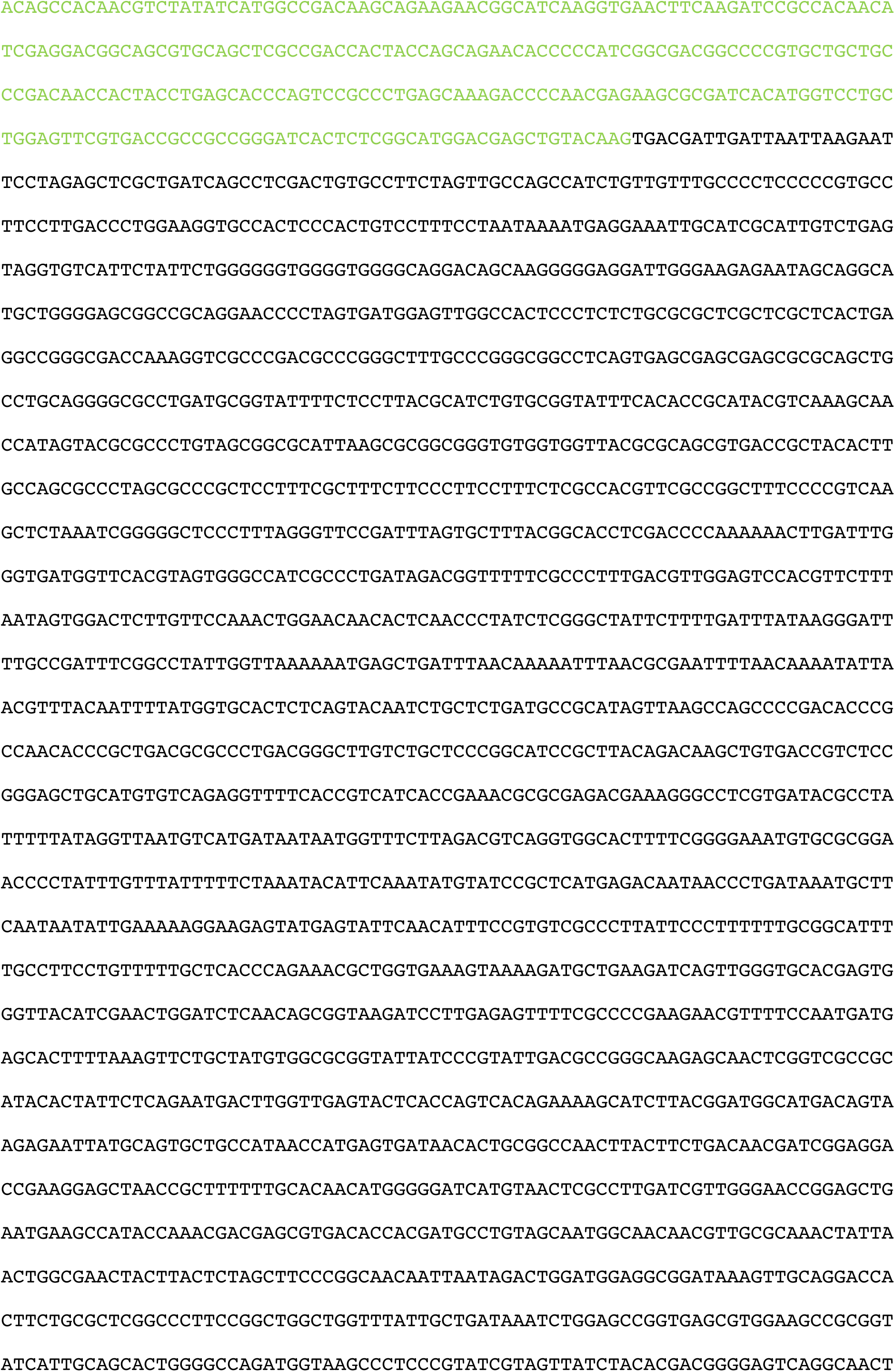
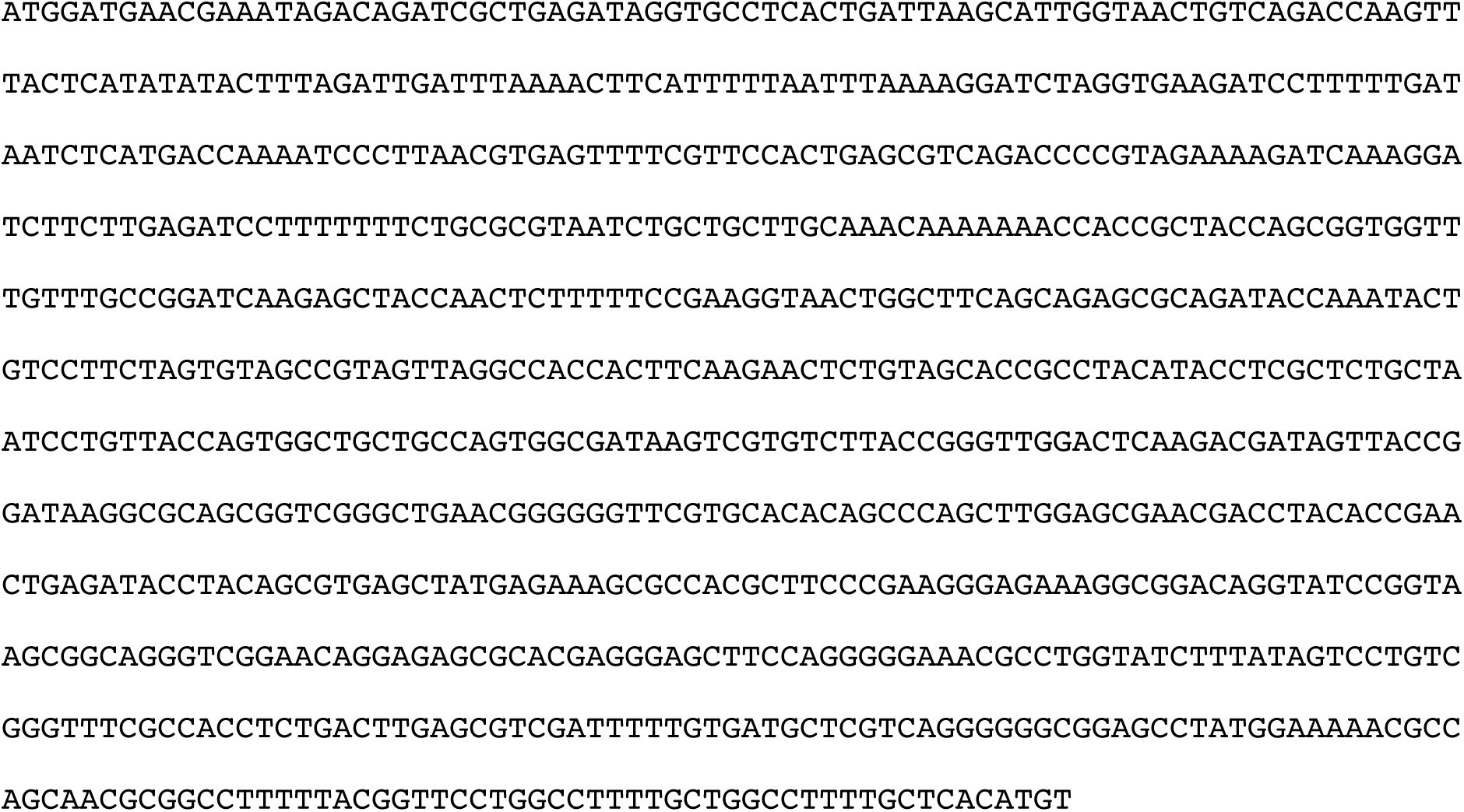

### Sequence of plasmid sgRNA-dCas9-2A-GFP

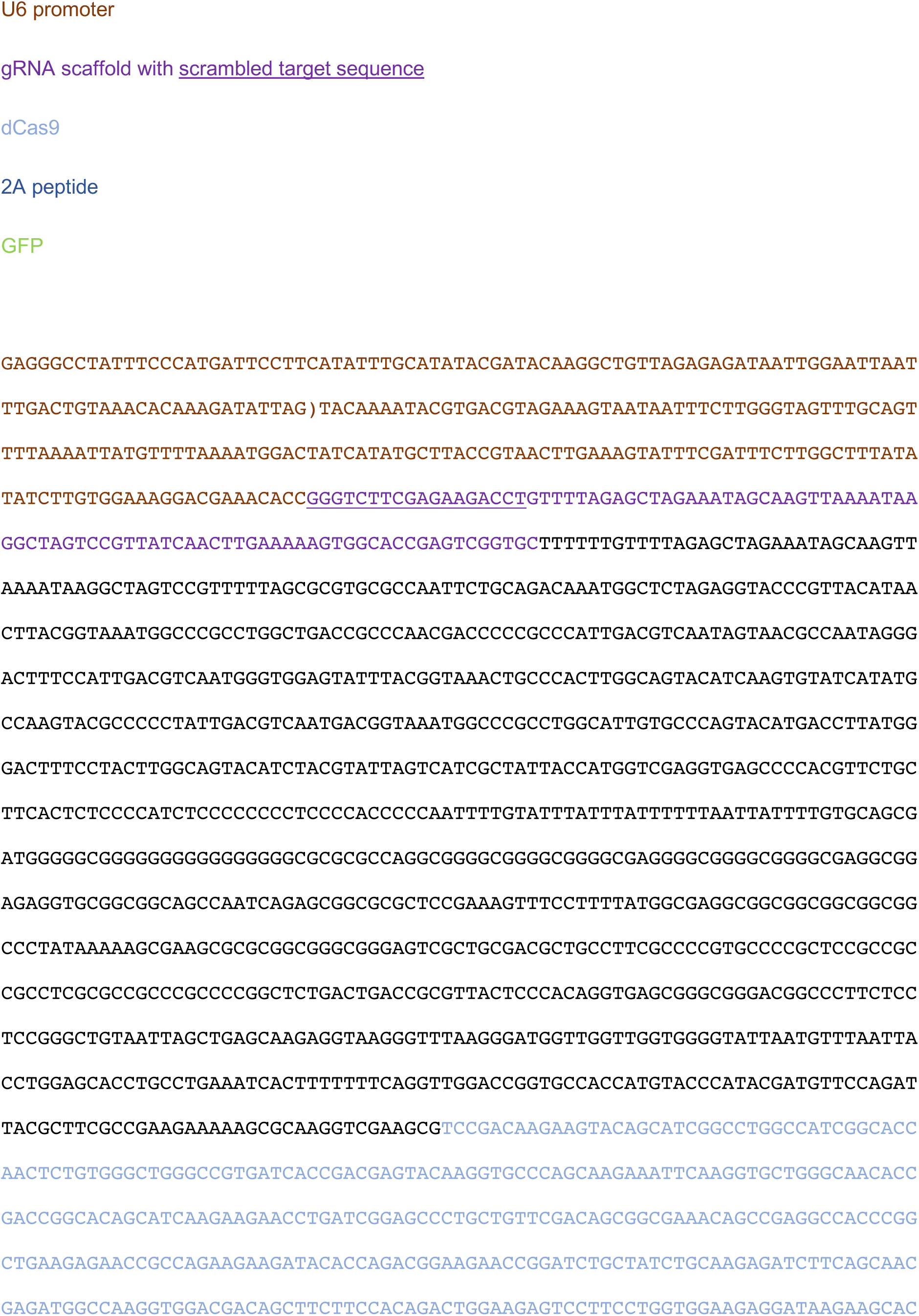
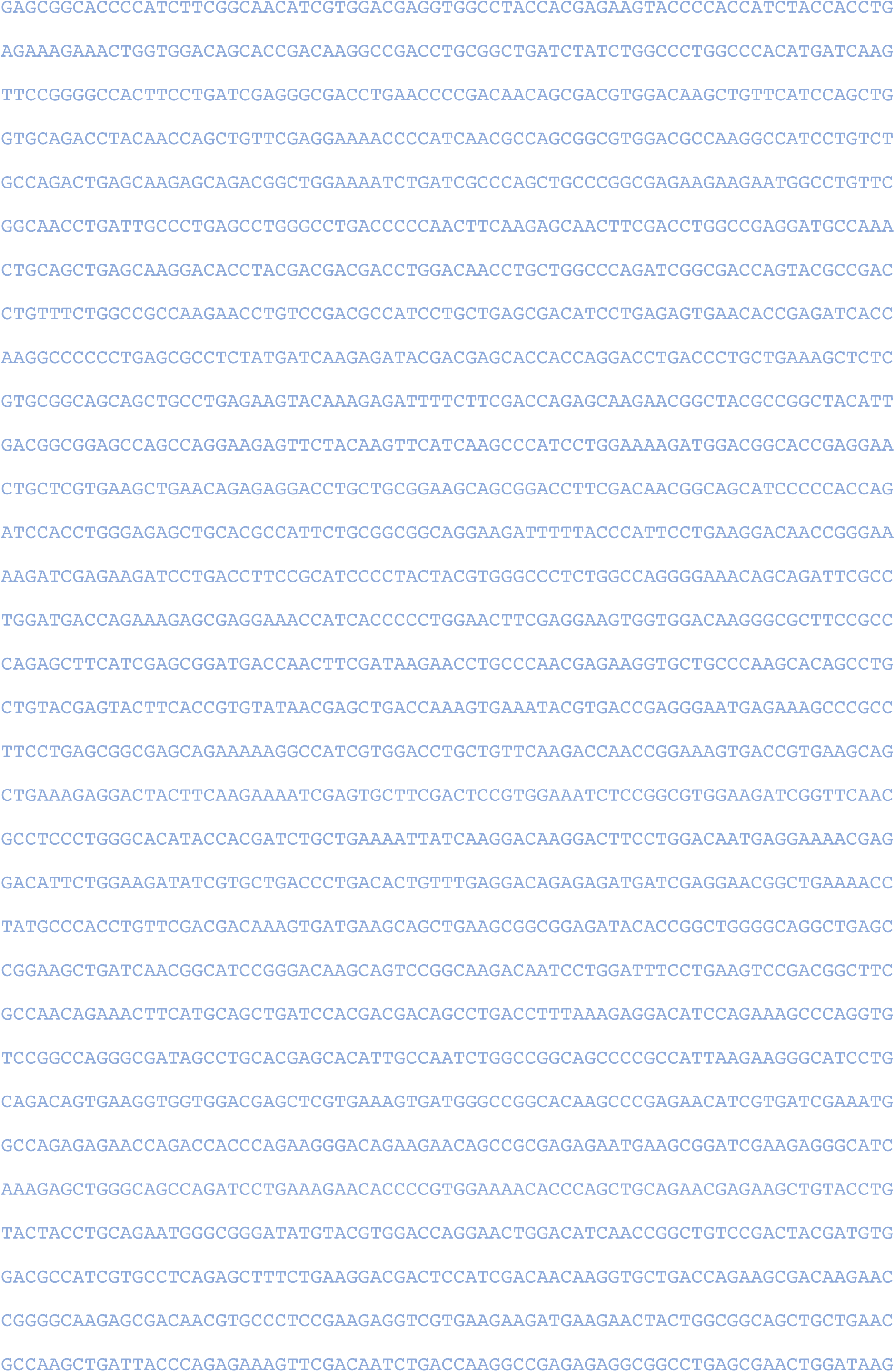
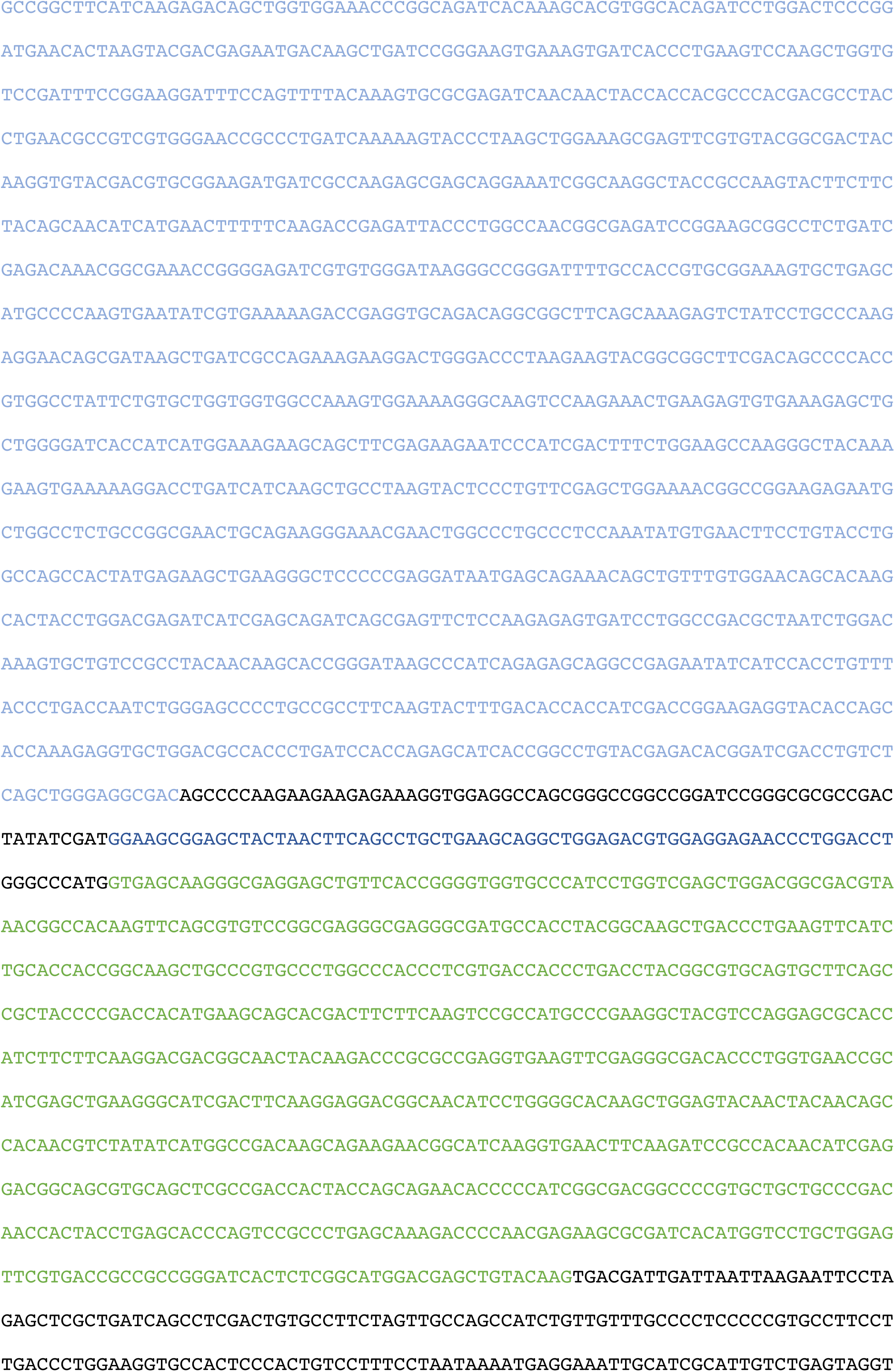
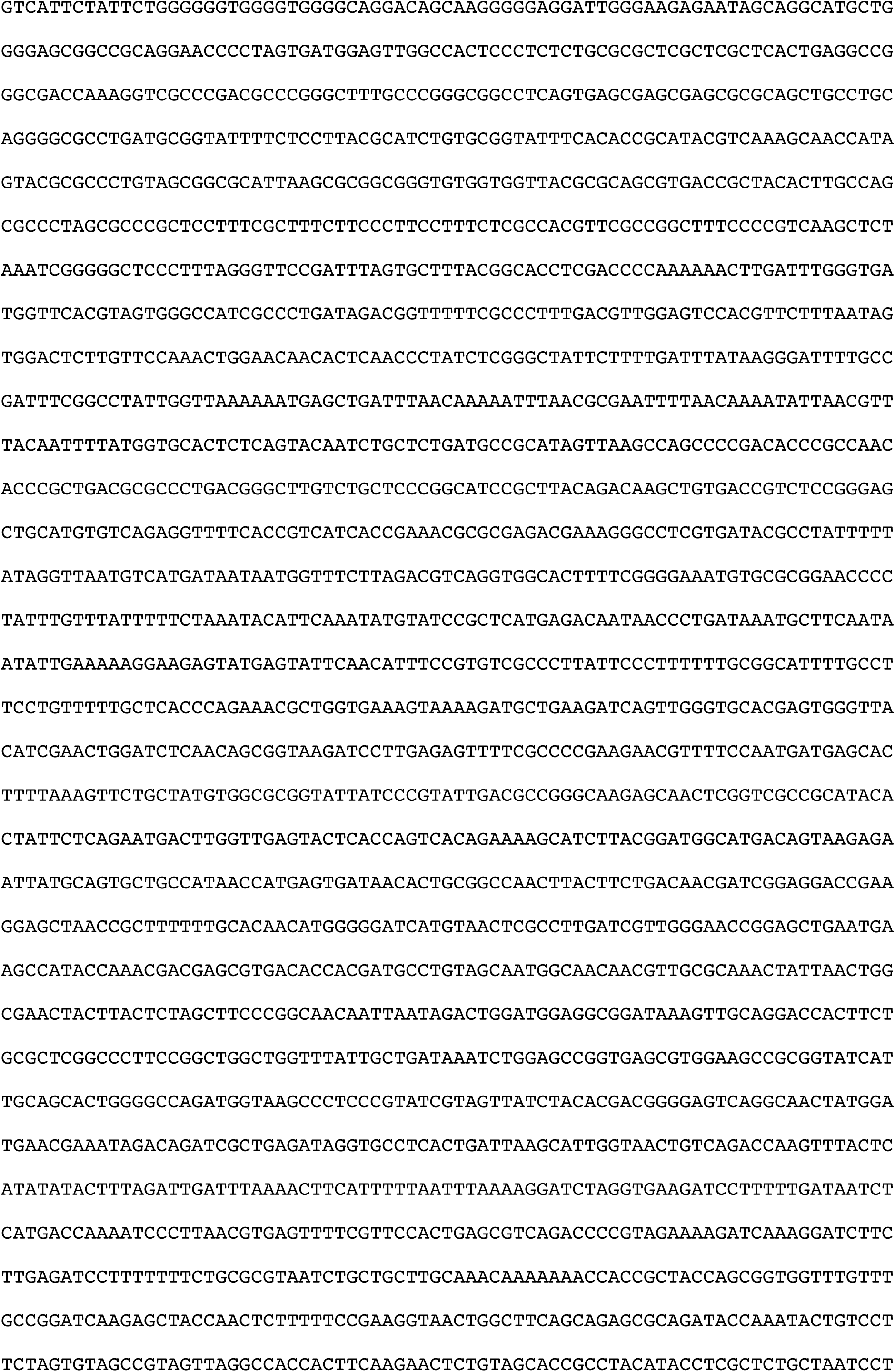
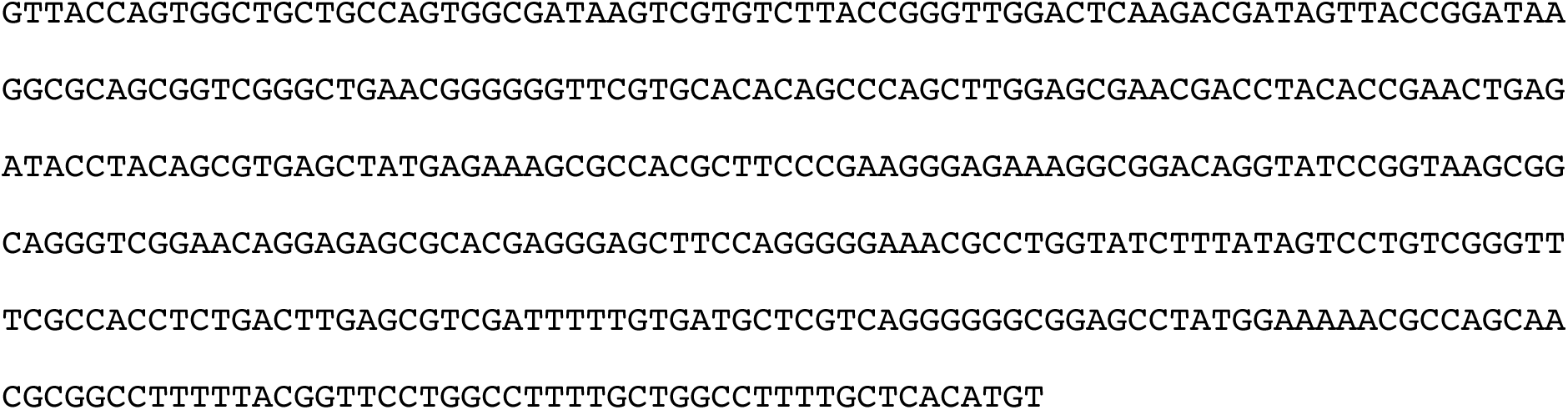

### Sequence of plasmid sgRNA-2A-GFP

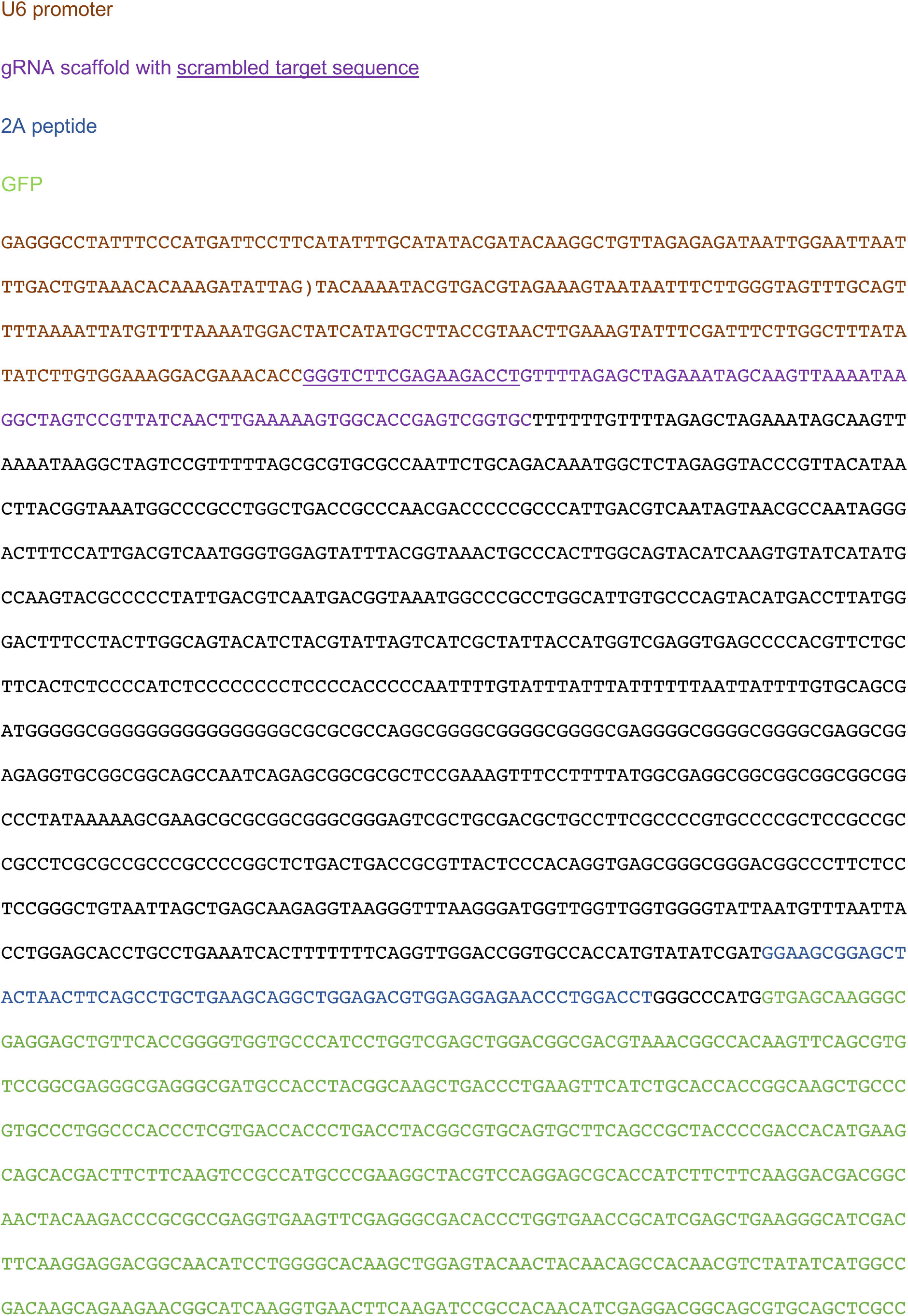
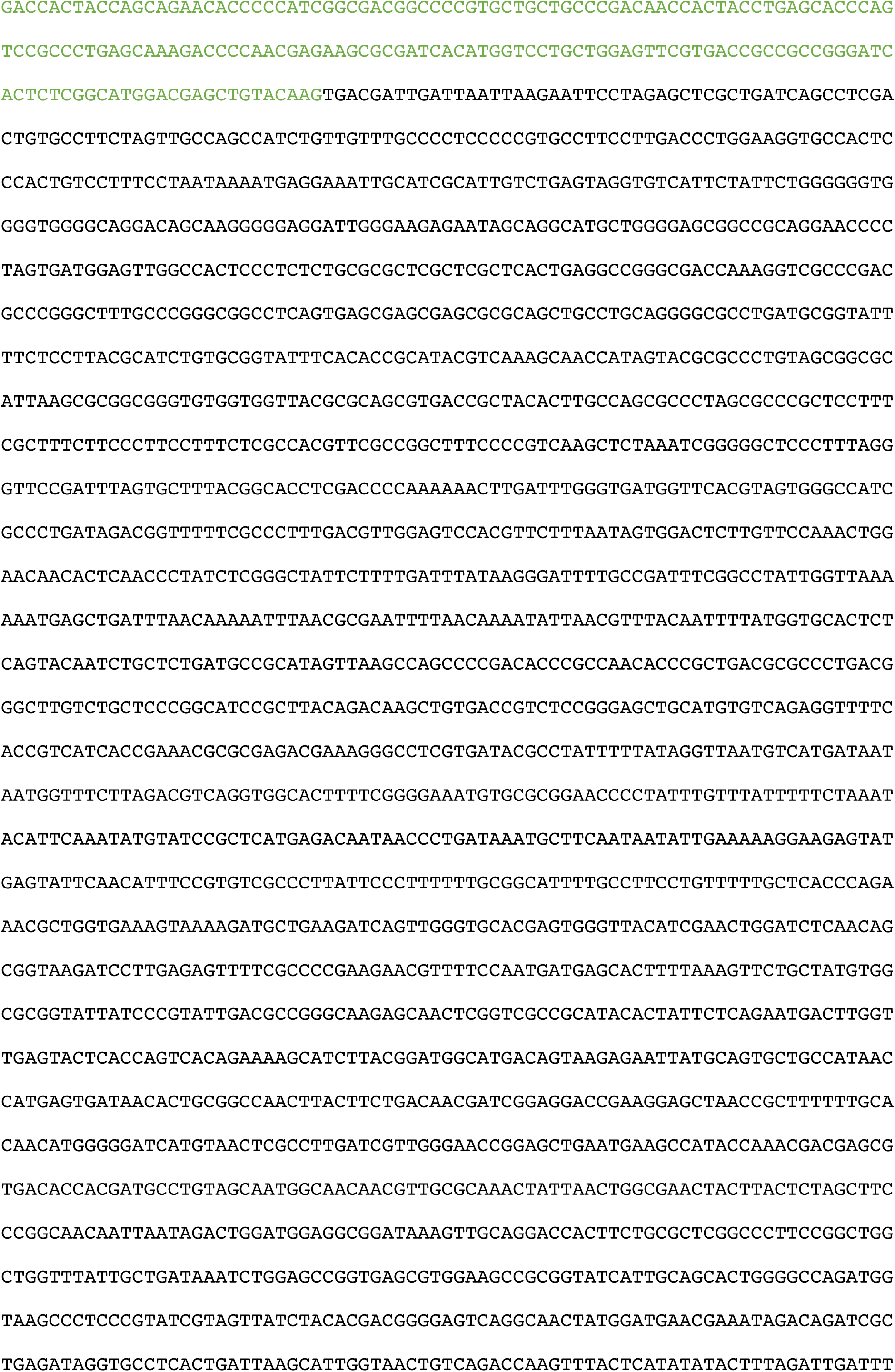
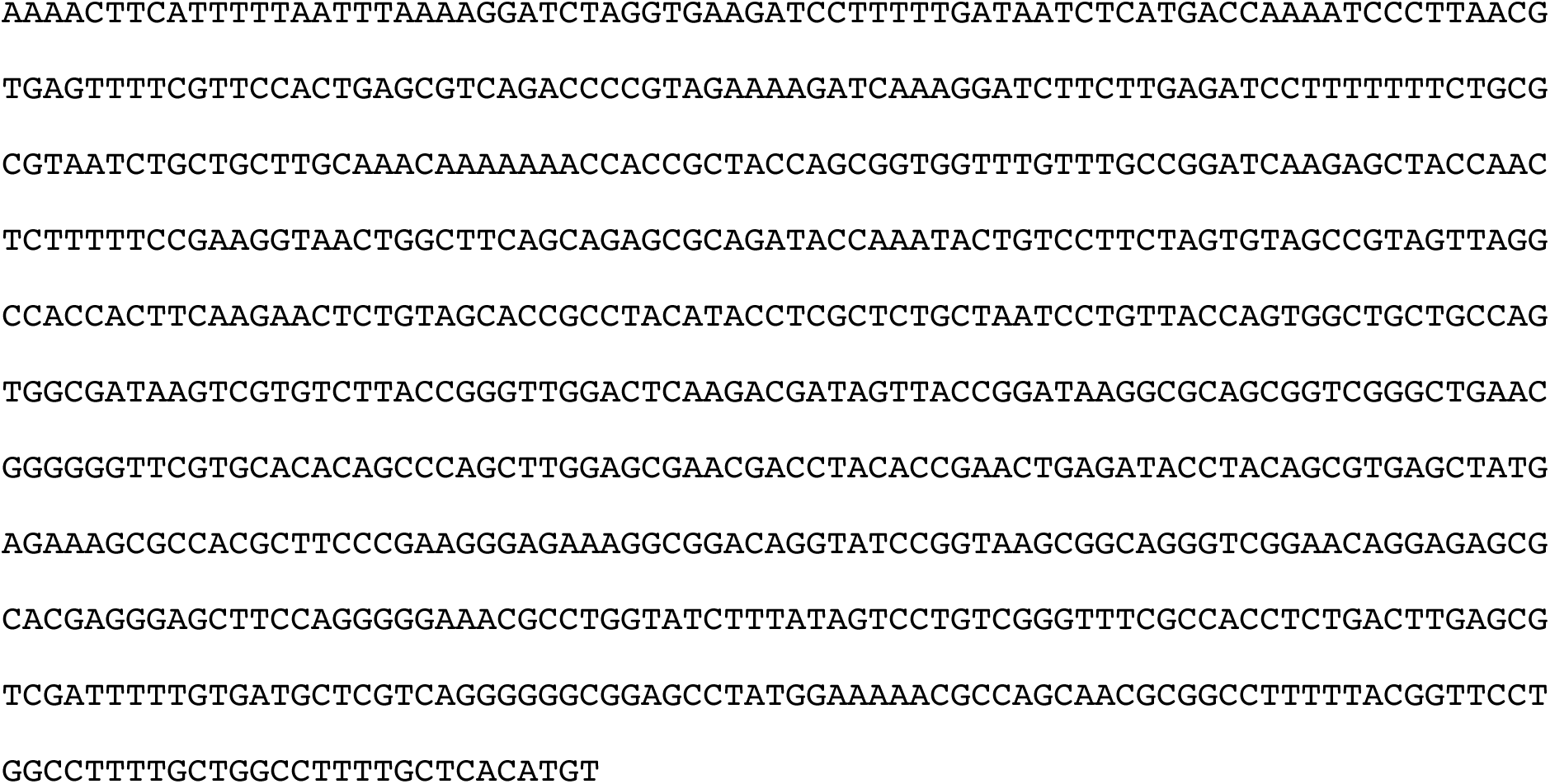

## Supplementary file 2 (Excel file)

### DAVID enrichment analysis

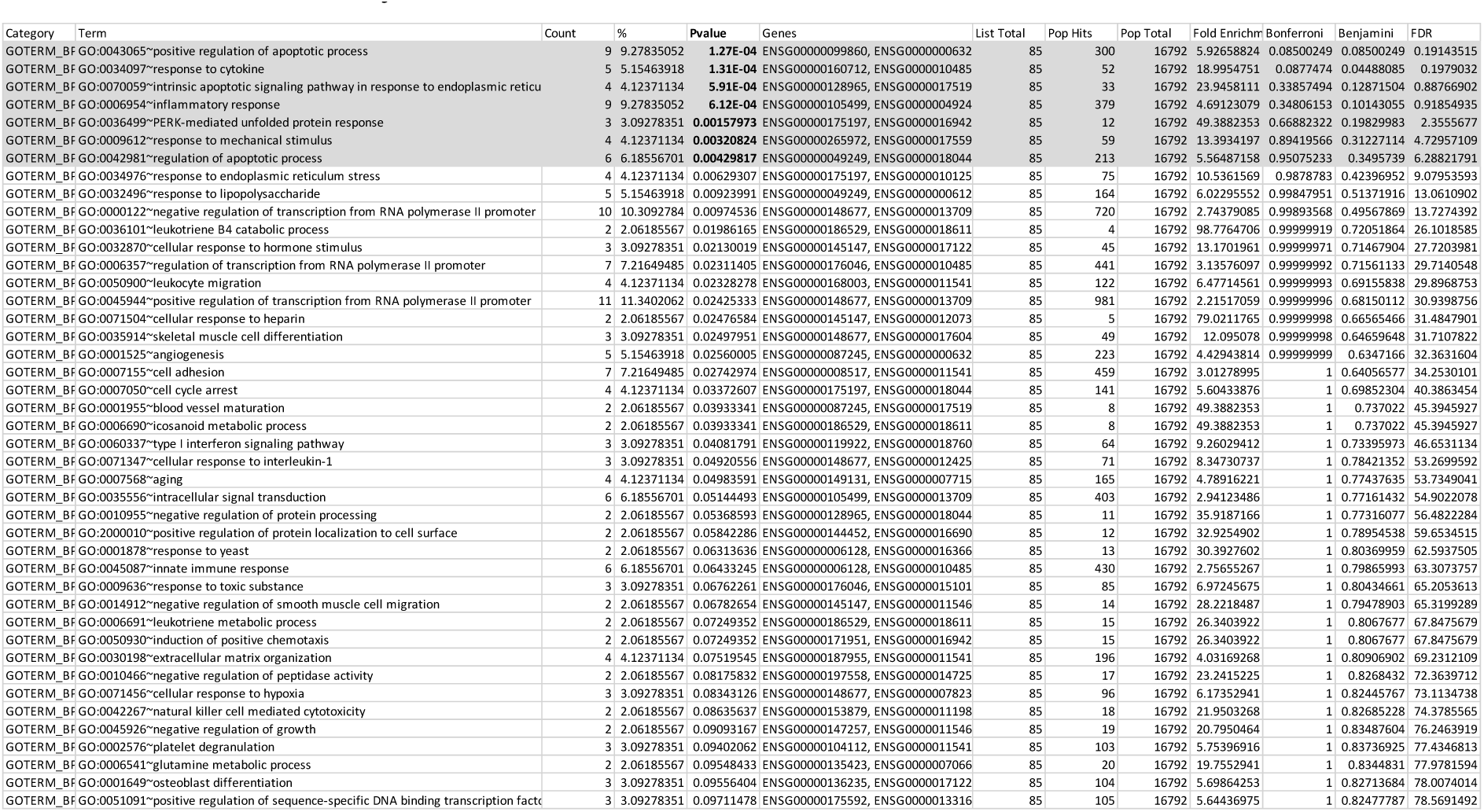

### Enriched pathways

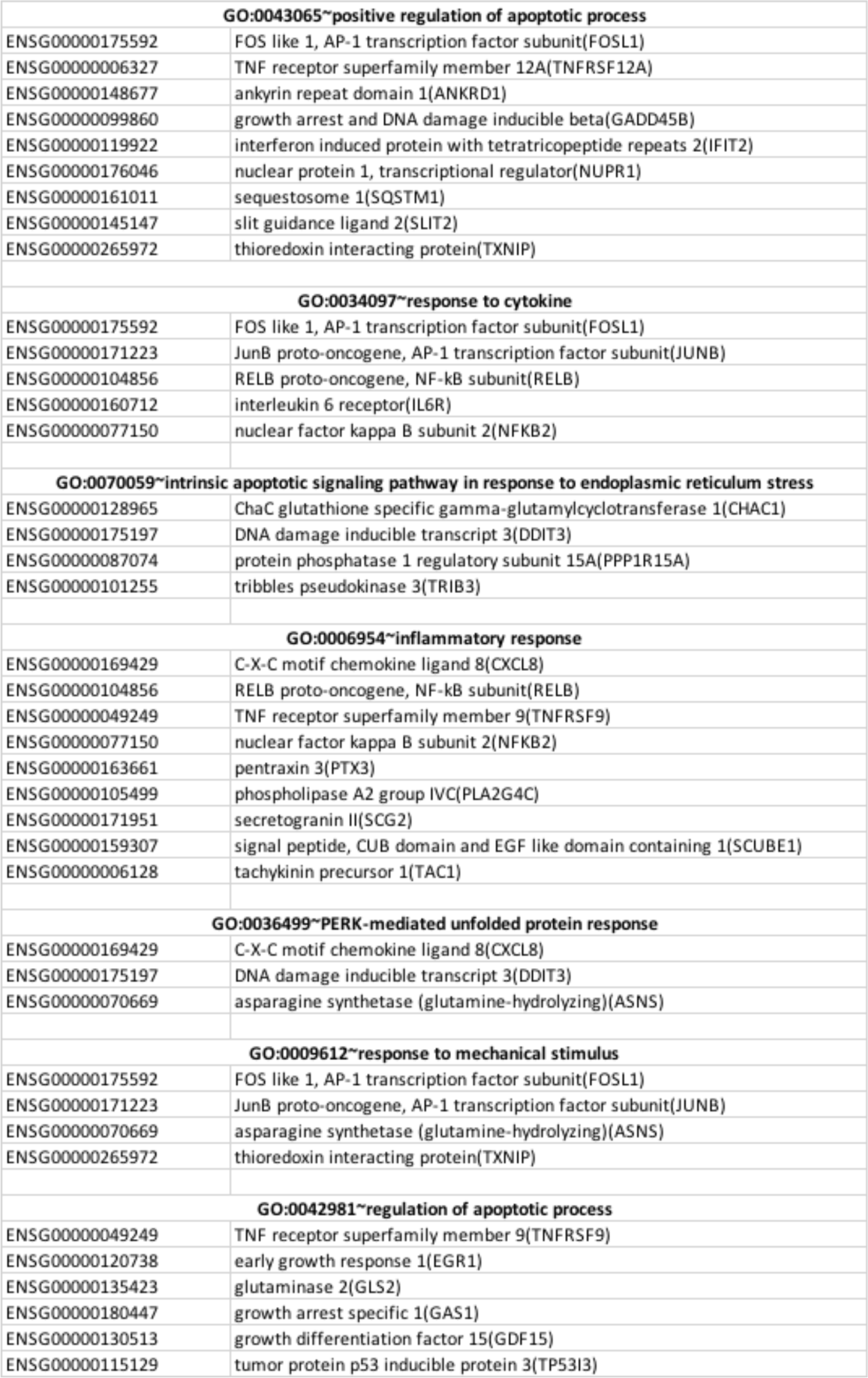

### David total 30 genes

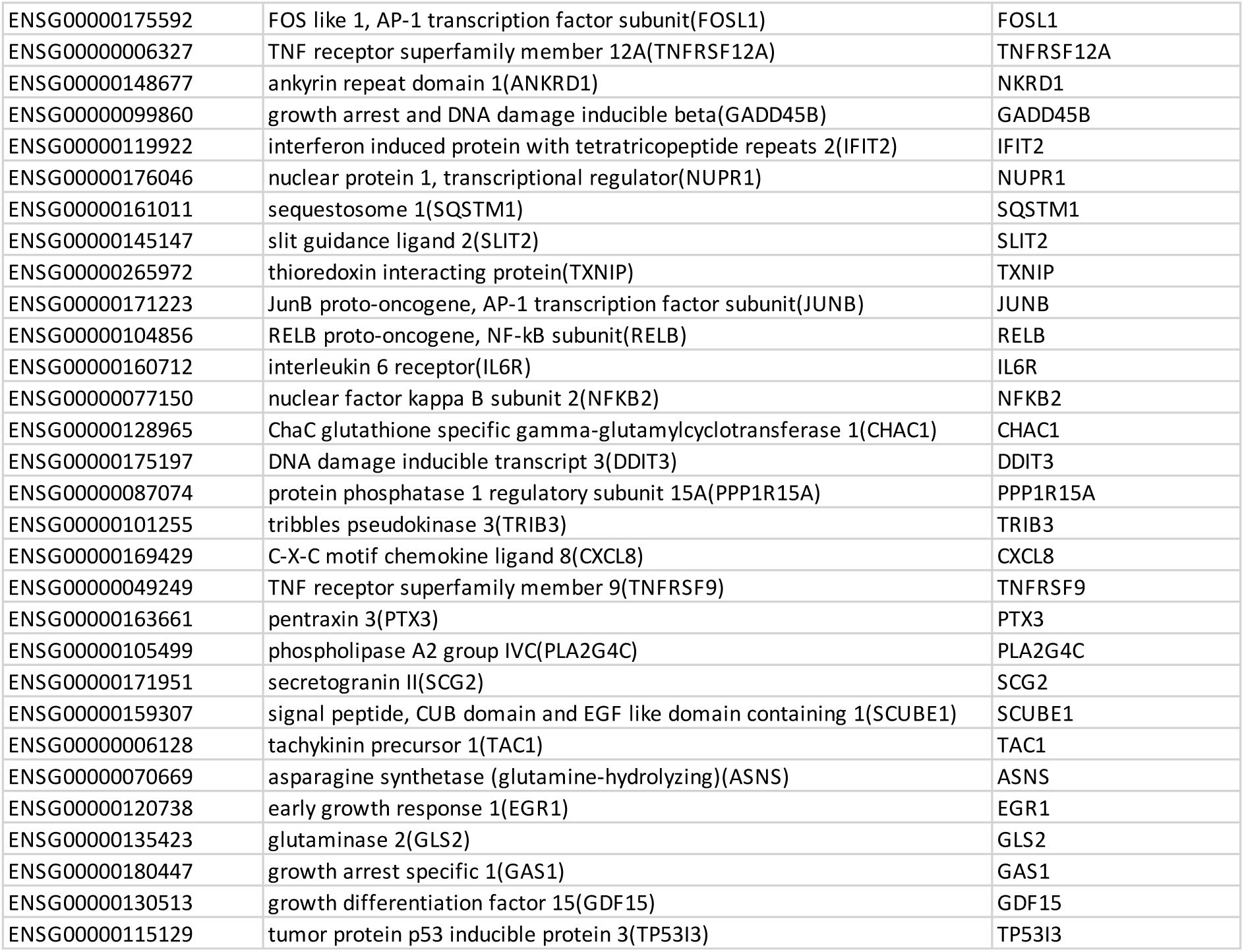

## Notes

https://www.ncbi.nlm.nih.gov/geo/query/acc.cgi?acc=GSE118277

